# Molnupiravir combined with different repurposed drugs further inhibits SARS-CoV-2 infection in human nasal epithelium *in vitro*

**DOI:** 10.1101/2022.01.10.475377

**Authors:** Hulda R. Jonsdottir, Denise Siegrist, Thomas Julien, Blandine Padey, Mendy Bouveret, Olivier Terrier, Andres Pizzorno, Song Huang, Kirandeep Samby, Timothy N.C. Wells, Bernadett Boda, Manuel Rosa-Calatrava, Olivier B. Engler, Samuel Constant

**Author notes:** These authors contributed equally to this work (Bernadett Boda, Manuel Rosa-Calatrava, Olivier B. Engler, Samuel Constant). Corresponding authors: Hulda R. Jonsdottir, Samuel Constant.

## Abstract

The severe acute respiratory syndrome coronavirus 2 (SARS-CoV-2), first identified in late 2019, has caused a worldwide pandemic with unprecedented economic and societal impact. Currently, several vaccines are available, and multitudes of antiviral treatments have been proposed and tested. Although many of the vaccines show high clinical efficacy, they are not equally accessible worldwide. Additionally, due to the continuous emergence of new virus variants, and generally short duration of immunity, the development of safe and effective antiviral treatments remains of the utmost importance. Since the emergence of SARS-CoV-2, substantial efforts have been undertaken to repurpose existing and approved drugs for accelerated clinical testing and potential emergency use authorizations. However, drug-repurposing using high throughput screenings in cellular assays, often identify hits that later prove ineffective in clinical studies. Our approach was to evaluate the activity of compounds that have either been tested clinically or already undergone extensive preclinical profiling, using a standardized *in vitro* model of human nasal epithelium. Secondly, we evaluated drug combinations using sub-maximal doses of each active single compound. Here, we report the antiviral effects of 95 single compounds and 30 combinations. The data show that selected drug combinations including 10 μM of molnupiravir, a viral RNA-dependent RNA polymerase (RdRp) inhibitor, effectively inhibit SARS-CoV-2 replication. This indicates that such combinations are worthy of further evaluation as potential treatment strategies against coronavirus disease 2019 (COVID-19).

## Introduction

In December 2019, a novel coronavirus (CoV) causing respiratory illness in humans was identified in Wuhan, China (1). In the following months, this virus, designated severe acute respiratory syndrome coronavirus 2 (SARS-CoV-2) (2), spread to every continent and currently there have been over 290 million documented cases worldwide and over 5 million deaths (3). The corresponding illness, termed coronavirus disease 2019 (COVID-19), presents with a range of symptoms and severity (4-8). The acute respiratory disease syndrome (ARDS) observed in severe cases of COVID-19 is driven by the cytokine storm, characterized by uncontrolled overproduction of soluble immune mediators. This results in sustained inflammation and tissue injury that can lead to low oxygenation and eventual death (9, 10). A recurring pattern of upregulation of pro-inflammatory cytokines, including IL-6, IL-1β, and TNF-α has been observed (11, 12) with additional secretion of various chemokines (MCP-1, IP-10, RANTES) resulting in the influx of immune cells into the pulmonary space (13-16). Despite the rapid development and current availability of several vaccines, effective and safe treatments against COVID-19 are still of extreme importance due to incomplete vaccine uptake, incomplete protective response, and the relatively short duration of immunity. Additionally, those who either cannot be vaccinated or choose not to be, are not protected against severe COVID-19. Development of new treatments traditionally takes many years; however, the rapid spread of the virus means that new treatment options against SARS-CoV-2 are urgently needed. Drug repurposing redirects existing or previously approved drugs as new therapeutics for new clinical indications in a time- and cost-effective manner. For example, remdesivir (also known as GS-5734) an intravenous ProTide prodrug (17) targeting the RNA-dependent RNA polymerase (RdRp) of Hepatitis C virus, was repurposed and conditionally approved as emergency treatment for COVID-19 early in the pandemic (18). However, due to the complex interplay between the host immune system and virus replication, drugs inhibiting viral replication alone are often not sufficient to fight the systemic viral infection and resulting sequelae. For early ambulatory treatment of COVID-19, various treatments combining compounds targeting viral replication with other drugs have been proposed to maximize therapeutic potential (19-21). In addition, experiences gained through the development of therapeutic strategies against other RNA viruses (e.g., HIV, HCV) underline the importance of combining several drugs targeting different viral proteins and/or host factors to prevent the emergence of drug resistant mutants (22-24). Due to the heterogeneity of *in vitro* cell lines used in different studies, direct comparison of drug efficacy is difficult. To allow better comparison of compounds, we developed a standardized *in vitro* screening model, using fully differentiated primary human nasal epithelium (MucilAir™). Applying a standardized test protocol, we re-evaluated a collection of compounds from the Medicines for Malaria Venture (MVV) open access collections (25) and the Calibr ReFRAME drug-repurposing library (26). In the current study, we assessed the antiviral potential of 99 single compounds at two study sites with a focus on orally active drugs. Subsequently, we selected molnupiravir, an orally bioactive RdRp inhibitor recently approved by the American Food and Drug Administration (FDA), as a base compound for combination therapy (27). We evaluated various concentrations of 30 drugs in combination with 10 μM molnupiravir and found that selected combinations have the potential to boost antiviral activity compared to single treatments, indicating a benefit of combined antiviral treatment against SARS-CoV-2.

## Materials and Methods

### Reconstituted nasal epithelium - MucilAir^™^

Human nasal epithelial cells were obtained from patients undergoing polypectomy, experimental procedures were explained in full, and all subjects provided informed consent. The study was conducted according to the declaration of Helsinki (Hong Kong amendment, 1989), and received approval from local ethics commissions. Cells were isolated from primary tissue as previously described (28) and expanded once (p1). Pooled nasal epithelial cells from 14 individual donors were then seeded on 6.5-mm Transwell^®^ inserts (cat #3470, Corning Incorporated, Oxyphen, Wetzikon, Switzerland) in MucilAir™ culture medium (EP04MM, Epithelix Sàrl, Geneva, Switzerland). Once confluent, air-liquid interface (ALI) was established and maintained for at least 28 days for mucociliary differentiation. Average culture time post ALI was 43 days.

### Compound selection and toxicity

Compounds were sourced from Calibr (California Institute for Biomedical Research, La Jolla, California, USA) or from MMV except for IFN-α (IF007 #3308269), IFN-λ (SRP3060), ebselen (E3520), dalbavancin (SML-2378) and nanchangmycin (SML2251 #0000040357) which were purchased from Sigma Aldrich (Buchs SG, Switzerland), while IFN-β (AF-300-02B) was sourced from Peprotech (LubioScience GmbH, Zürich, Switzerland). Compounds were selected using available data from primary screens using various cell lines (Vero E6, HeLa-ACE2, Calu-3) and organoids and were obtained blinded as 10 mM stock solution in dimethyl sulfoxide (DMSO). Compound toxicity (n=104) was evaluated by assessing the release of lactate dehydrogenase (LDH, Assay Kit-WST CK12-20, Dojindo EU, Munich, Germany) from damaged cells at 48 and 72h post treatment. Compound-containing cell culture media was replenished every 24h. Trans-epithelial electrical resistance (TEER) was monitored at 24, 48, and 72h, while ciliary beating frequency (CBF) was assessed at 72h by capturing 256 images at a high frequency rate (125 frames per second) at room temperature (RT). CBF was calculated using Cilia-X software (Epithelix Sàrl, Geneva, Switzerland).

### Determination of epithelial integrity

Epithelial integrity was determined by quantifying TEER at 48 and 72h post infection (hpi) using the EVOM3/EVOMX Volt/Ohm Meter and the accompanying chopstick electrode (STX2/STX2-plus, World Precision Instruments, Friedberg, Germany). 200 μL OptiMEM (Gibco, Thermo Fisher, Basel, Switzerland) or 0.9% NaCl solution was added to the apical compartment prior to measurement. Total resistance values (Ω) were converted to TEER (Ω.cm^2^) by subtracting 100 Ω for the resistance of the polyester membrane and multiplying by 0.33 cm^2^.

### Virus propagation

SARS-CoV-2 (reference strain: BetaCoV/France/IDF0571/2020, EPI_ISL_406596) used at the primary testing site was acquired from EVAg (Emerging Viral Diseases Medical Faculty, Marseille, France) and propagated on Vero E6 cells (provided by Prof. Dr. Volker Thiel, University of Bern, Bern, Switzerland) in Minimum Essential Medium (MEM; Seraglob, Schaffhausen, Switzerland) supplemented with 2% Fetal Bovine Serum (FBS; Seraglob) at 37°C, 5% CO2 and >85% relative humidity (rH) for 3 days prior to harvest. The SARS-CoV-2 strain used at the second testing site was isolated from a patient in a French clinical cohort of patients with COVID-19 (NCT04262921) in January 2020 at the Department of Infectious and Tropical Diseases, Bichat Claude Bernard Hospital, Paris (BetaCoV/France/IDF0571/2020, EPI_ISL_411218) (29). Tissue culture infectious dose 50 (TCID50/ml) was determined by a limiting dilution assay under the same culture conditions with the Spearman-Kärber method (30, 31) or the Reed & Muench statistical method (32, 33), depending on testing site.

### Virus infection

Prior to infection, MucilAir™ reconstituted nasal epithelium, was washed twice with OptiMEM (Gibco) warmed to 37°C and basal media replenished with warm MucilAir™ cell culture media. 5×10^4^ TCID_50_ of SARS-CoV-2 (for a theoretical MOI of 0.1) were added to the apical compartment diluted in OptiMEM and incubated at 37°C, 5% CO_2_ and >85% rH for 1h. Mock controls were exposed to the same volume of OptiMEM only. Subsequently, virus inoculum was removed, and the epithelium transferred to cell culture media containing test or control compounds. Apical virus release was assessed at 48 and 72 hpi by washing the apical side with 200 μl OptiMEM for 10 min at 37°C. During harvest of viral particles from the apical compartment, while the epithelium is submerged, TEER was assessed as described above. Test and control compounds were replenished every 24 h for the duration of the experiment (72h). Remdesivir was applied as a positive control at a concentration of 5 μM (Gilead Sciences Inc., Foster City, California, USA or MedChemExpress, (HY-104077)). All compounds were diluted in DMSO and final concentration of DMSO in the antiviral assay was standardized as 0.3% for single compounds and 0.4% for combinations. Experimental layout is presented in Figure 1.

**Figure 1:**
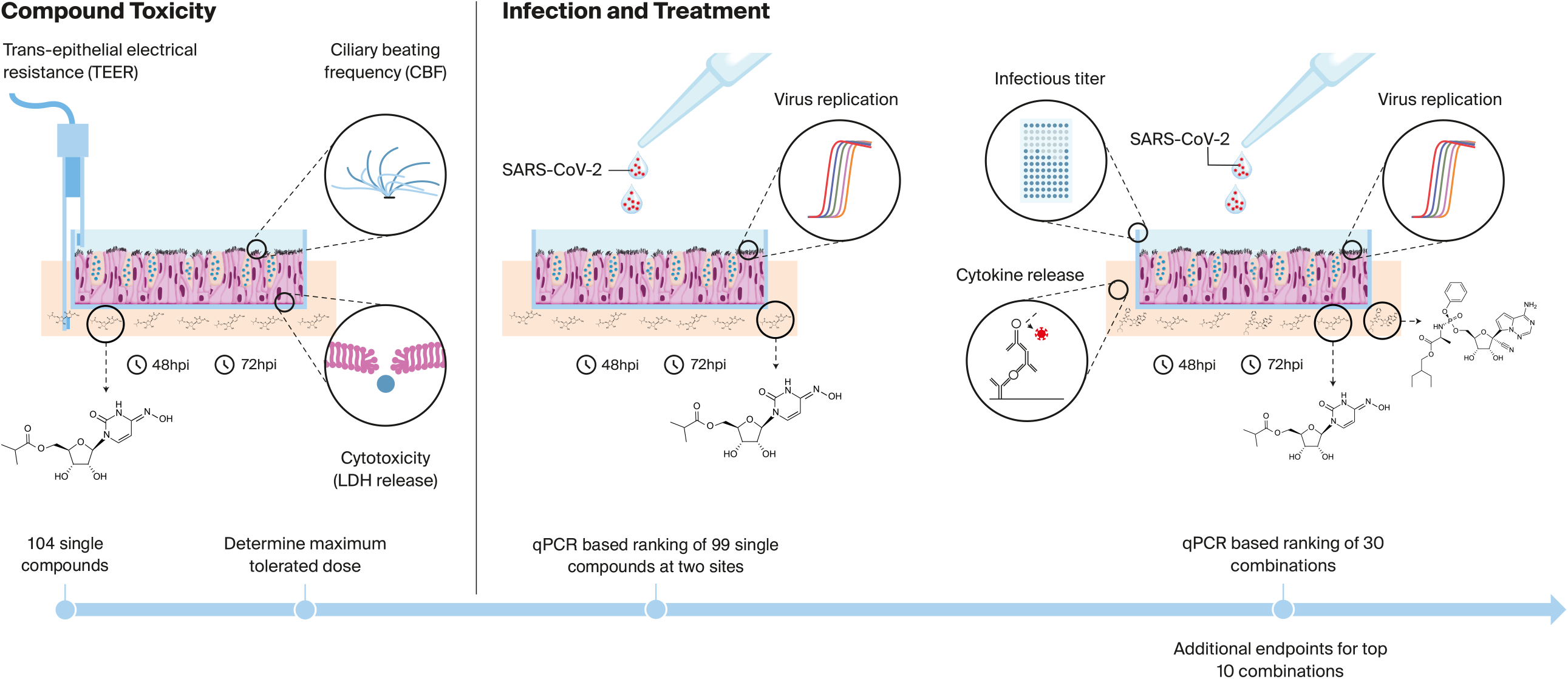
Overview over experimental design and layout. Toxicity of 104 single compounds was determined in reconstituted *in vitro* nasal epithelium in order to determine the maximum tolerated dose (≤30 μM). Compounds were applied basally, mimicking systemic administration. Three end-points were assessed, trans-epithelial electrical resistance (TEER), cytotoxicity, and the ciliary beating frequency of ciliated cells. Subsequently, 99 single compounds were screened for antiviral activity against SARS-CoV-2. Additionally, 30 combinations based on the orally bioactive RdRp inhibitor molnupiravir were tested for antiviral activity with additional parameters (infectious virus and cytokine secretion) determined for top candidates.

### RNA extraction

200 μl of apical wash was harvested at 48 and 72 hpi and 100 μl inactivated in 400 l AVL buffer (Qiagen, Hombrechtikon, Switzerland) for 10 min at RT then 400 μl of absolute ethanol were added to each sample. Viral RNA was extracted from 500 μl with the MagNaPure 96 system (Roche, Basel, Switzerland) according to the manufacturer’s instructions and extracted RNA eluted in 100 μl. Alternatively, 140 μl were used for viral RNA extraction with the QIAamp^®^ Viral RNA kit (Qiagen, Hombrechtikon, Switzerland), obtaining 60 μl of eluted RNA. At the end of each experiment, epithelia were lysed in Qiazol (Qiagen, Hombrechtikon, Switzerland) for a total of 20 min at RT. RNA was extracted using the Qiagen RNeasy Plus Universal Mini kit according to the manufacturers’ instruction, eluted in 30 μl of nuclease free water, and diluted 1:10 prior to gene expression analysis.

### qRT-PCR

Viral RNA in apical secretions was quantified using the TaqMan™ Fast-Virus-1 Step master mix (Thermo Fisher, Basel, Switzerland) with these cycling parameters: 50°C for 1 min, 95°C for 20 s, 45 cycles of 95°C for 3 s and 60°C for 30 s using the LightCycler 96 system (Roche, Basel, Switzerland) or the StepOnePlus™ Real-Time PCR System (Applied Biosystems, Massachusetts, USA) with the following primers against SARS-CoV-2 non-structural protein 14 (nsp14): Fwd: 5’-TGGGGYTTTACRGGTAACCT-3’, Rev: 5’-AACRCGCTTAACAAAGCACTC-3’. Probe: 5’-FAM-TAGTTGTGATGCWATCATGACTAG-TAMRA-3’ (protocol by Leo Poon, Daniel Chu, and Malik Peiris; School of Public Health, The University of Hong Kong, Hong Kong). Changes in gene expression were calculated using the 2^-ΔCt^ method and reported as the fold reduction over infected vehicle treated inserts (virus control, 0.3-0.4% DMSO). Intracellular RNA was quantified using the SuperScript III Platinum One-step SYBR Green master mix (Thermo Fisher, Basel, Switzerland) and these cycling parameters: 50°C for 1 min, 95°C for 5 min, 50 cycles of 95°C for 15 s, 60°C for 30 s (53°C for nsp14), 40°C for 1 min followed by a melting curve to confirm product specificity or the StepOnePlus™ Real-Time PCR System (Applied Biosystems) using the previously mentioned primers against nsp14 and the following primers against GAPDH: Fwd: 5’-GAAGGTGAAGGTCGGAGTCAAC-3’, Rev: 5’-CAGAGTTAAAAGCAGCCCTGGT-3’ (34). Changes in intracellular gene expression were calculated using the 2^-ΔΔCt^ method and reported as the fold reduction over infected vehicle treated inserts (virus control, 0.3-0.4% DMSO).

### Determination of infectious titer

To quantify infectious virus in apical secretions, apical wash samples (72 hpi) were diluted in MEM supplemented with 2% FBS (Seraglob) and titrated as described above. Vero E6 cells modified to constitutively express the serine protease TMPRSS2 (35) were obtained from the Centre for AIDS Reagents (National Institute for Biological Standards and Control) and were incubated for 3 days at 37°C, 5% CO_2_ and >85% rH prior to CPE determination by crystal violet staining. Infectious titer was determined with the Spearman-Kärber method (30, 31, 33) and reported as tissue culture infectious dose 50 per ml (TCID_50_/ml).

### Determination of cytokine release

Secretion of CXC motif chemokine ligand 10 (CXCL10/IP-10, DY266) and CC chemokine ligand 5 (CCL5/RANTES, DY278) were quantified by DuoSet^®^ ELISA (R&D systems, Bio-Techne GmbH, Wiesbaden, Germany) under BSL-3 conditions according to the manufacturer’s instructions using the appropriate ancillary reagent kit (DY007).

### Statistical analysis

Statistical significance between single molnupiravir treatments and combinations were determined by ordinary one-way analysis of variance (ANOVA) with Dunnett’s multiple comparison test. Significance between single and combination treatment for other compounds was determined by ordinary one-way ANOVA with Sidak’s multiple comparison test. p-values <0.05 were considered statistically significant.

## Results

### Compound toxicity

Tolerated concentrations of selected compounds were evaluated up to 30 μM on pooled nasal epithelium (MucilAir™) by monitoring trans-epithelial electrical resistance (TEER), the release of lactate dehydrogenase (LDH), and ciliary beating frequency (CBF). Compounds applied in basal culture media were replenished every 24 h in accordance with our standardized protocol (29) and TEER measured on days 1, 2, and 3. The maximum tolerated dose (MTD ≤30 μM) for each of the tested compounds is reported in Table S1. Examples of a well-tolerated compound, molnupiravir, and a compound inducing toxic effects, LY2228820, are shown in Figure S1. A decrease in TEER is a clear indicator of cellular toxicity in this system and its measurement is compatible with virus replication and high biosafety laboratory constraints. Interestingly, when measuring TEER in infection/treatment experiments, some of the tested substances including osimertinib, midostaurine, ZLVG CHN2 and ozanimod, led to a loss of epithelial integrity (TEER<100 Ω.cm^2^) beyond that observed for vehicle control. Such compounds were excluded from further analysis.

### Antiviral activity of single compounds against SARS-CoV-2

We evaluated the antiviral potential of 99 single compounds against apical SARS-CoV-2 infection in re-constituted nasal epithelium. One hour post infection, epithelia were treated basally with different concentrations of the selected compounds. Antiviral activity was assessed 48 and 72 h post infection (hpi) by measuring viral RNA in apical wash (48 and 72 hpi) and intracellularly (72 hpi). Summarized data of all tested compounds can be found in Table S1 (n=95, 4 compounds were excluded based on the loss of tissue integrity). Of the 99 compounds tested, 16 exhibited >1 log reduction in viral RNA in either apical wash (Figure 2a) or intracellularly (Figure 2b) at 72 hpi at the highest concentration tested. In general, the reduction of viral RNA was equal or even more pronounced when measured intracellularly compared to apical wash. Those compounds exhibiting strong inhibition of the replication of SARS-CoV-2 in this study primarily target individual parts of the viral replication machinery. The most efficient inhibitor was PF-00835231, a SARS-CoV-2 specific protease inhibitor, resulting in a 4-5 log reduction of viral RNA at 30, 10 (Figure 2) and 5 μM (Figure 3). Following close behind are the RNA dependent RNA polymerase (RdRp) inhibitors remdesivir and molnupiravir, that also resulted in efficient reduction of SARS-CoV-2 replication at various concentrations. Remdesivir, included as positive control in all screening series, resulted in 3-4 log reduction in viral RNA in apical wash at 10, 5 (Figure 2) and 1.15 μM (Figure 3). Similar reduction (3.3 log) was observed after treatment with an orally active RdRp inhibitor, molnupiravir at 30 μM (Figure 2a). Both remdesivir and molnupiravir were tested multiple times at two separate testing sites with minimal deviation, demonstrating the robustness of our infection protocol, detection methods and assay transferability (Figure S2). Four other compounds, the HIV-specific protease inhibitors, nelfinavir and ritonavir as well as the protease inhibitors TO-195 and ONO-3307, resulted in an average 2-3 log reduction of SARS-CoV-2 RNA at the highest concentration tested (30 μM). Digitoxin, a cardiac glycoside, was also antiviral at a low concentration (0.36 μM) to similar levels as these non-specific protease inhibitors. Narasin, a coccidiostat and anti-bacterial also exhibited slight antiviral activity. Nafamostat, inhibitor of TMPRSS2 transmembrane serine protease, essential for SARS-CoV-2 host cell entry, resulted in average viral RNA reduction of 1.4 logs in apical wash at 30 μM, while reduction in intracellular RNA was four times higher. Furthermore, in our experimental model, treatment with emetine, ivermectin, homoharringtonine (HHT), alisporivir, camostat, and mitoguazine resulted in borderline one log reduction of viral RNA in either apical wash or intracellularly at the highest concentration tested (Figure 2).

**Figure 2:**
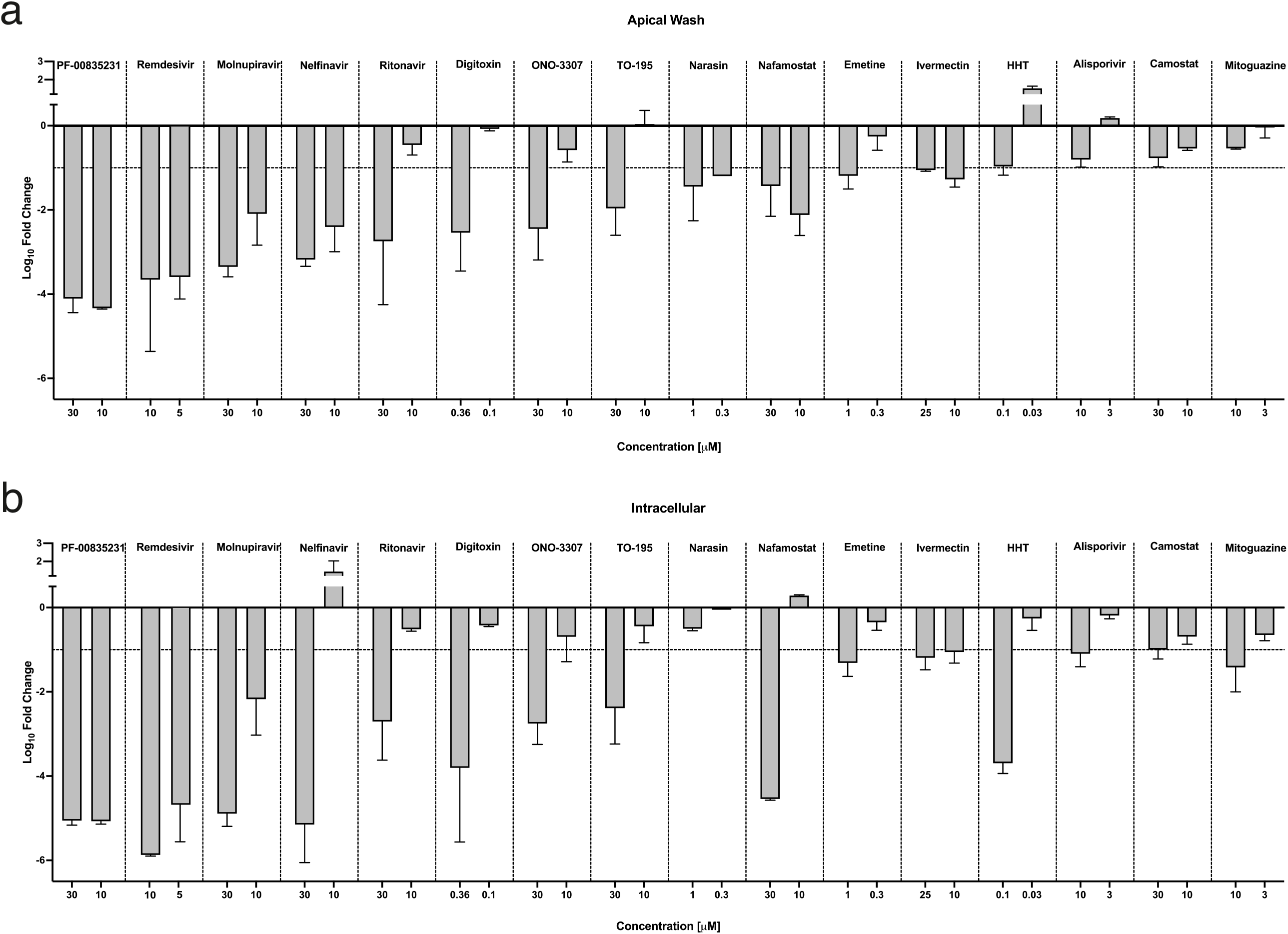
Antiviral activity of single compounds against SARS-CoV-2. Antiviral activity of single compounds as detected by qPCR in **a)** apical wash and **b)** intracellular lysate at 72 hpi. Top 16 compounds based on the viral content of the apical wash at 72 hpi (n=2-3, except for conditions tested at both investigatory sites: n=5 for 10 μM nelfinavir, camostat; n=6 for 30 μM molnupiravir; and n=12 for 10 μM remdesivir and molnupiravir. HHT: homoharringtonine. Data are represented as mean ± standard deviation (SD).

**Figure 3.**
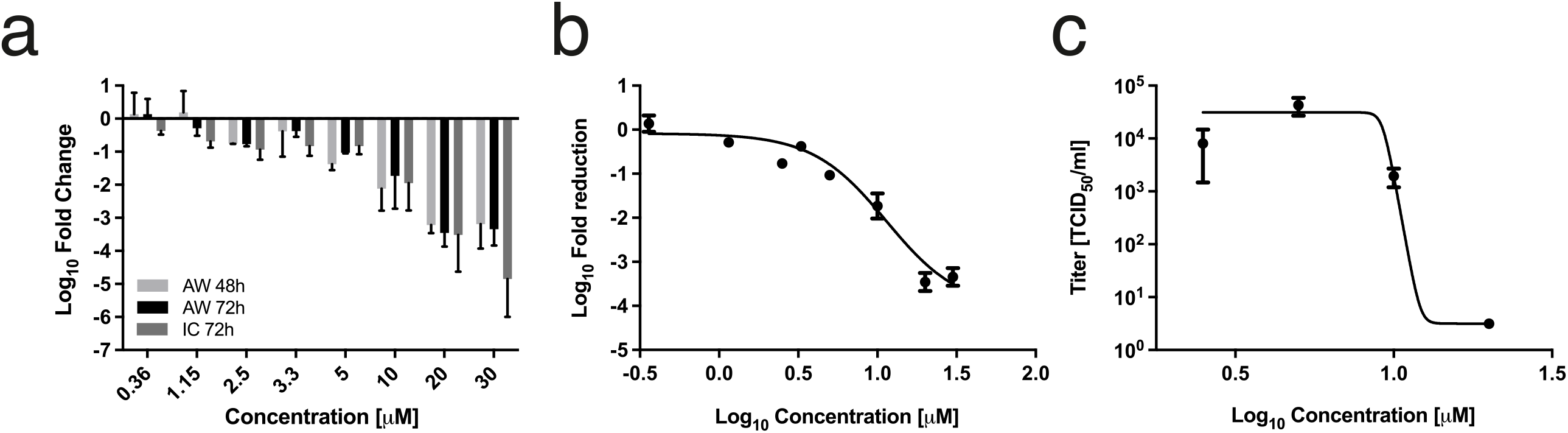
Determination of IC_50_ for Molnupiravir. **a)** Antiviral activity of molnupiravir at 48 and 72 hpi as detected by qPCR in apical wash. Data are represented as mean ± SEM, n=6, 6, 2, 6, 2, 12, 4 and 6 for 0.36, 1.15, 2.5, 3.3, 5, 10, 20 and 30 μM, respectively. **b)** qPCR dose response curve, IC_50_ determined at 11.1 μM. **c)** TCID_50_/ml dose response curve, IC_50_ determined at 9.2 μM (n=2-4).

**Figure 4:**
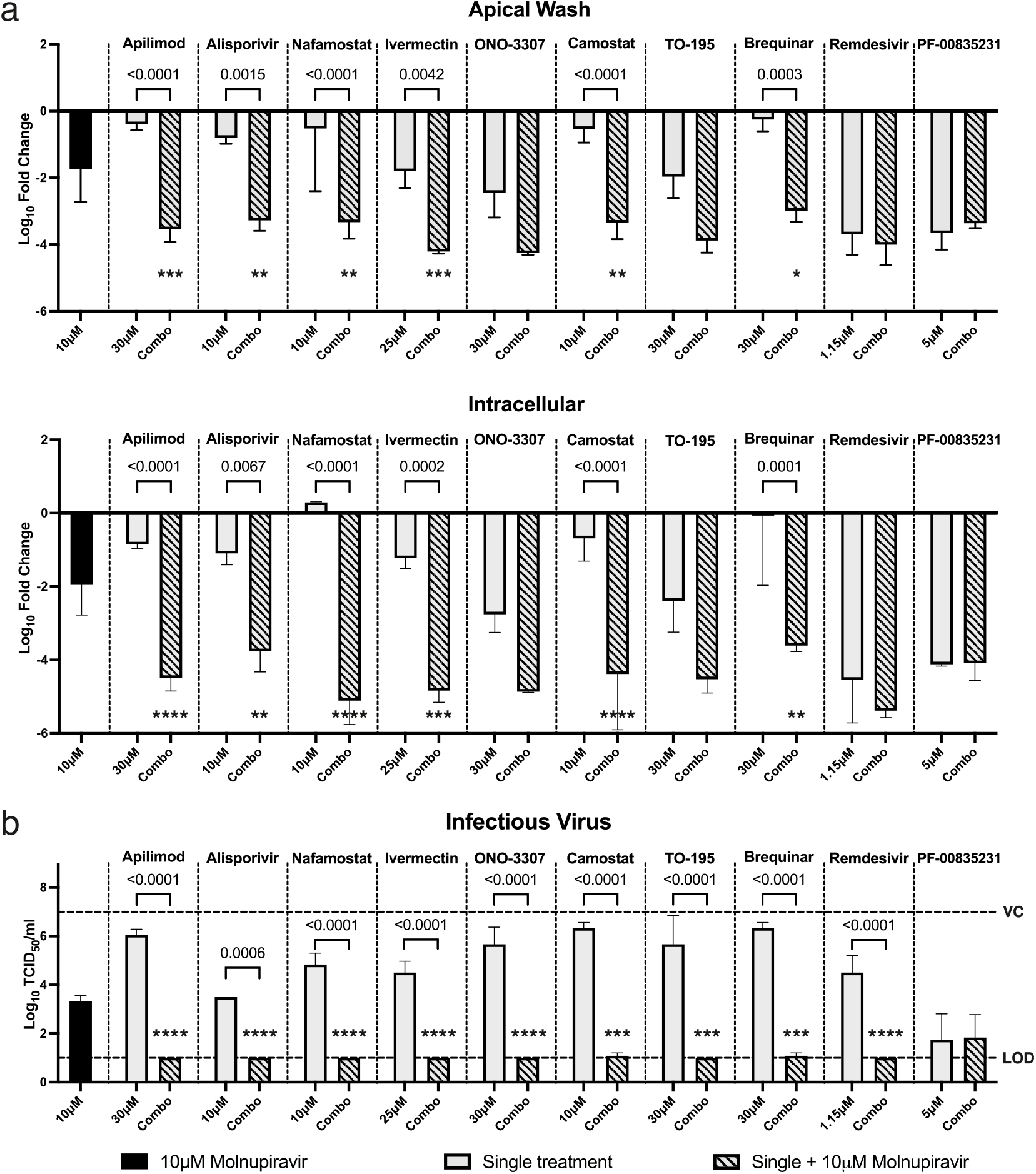
Antiviral activity of compound combinations against SARS-CoV-2. Antiviral activity of molnupiravir-based combinations against SARS-CoV-2 as determined by **a)** qPCR in apical wash or intracellularly at 72 hpi (n=2-4, and n=12 for molnupiravir) and **b)** TCID_50_/ml in apical wash at 72 hpi (n=2-4). Dashed lines represent the limit of detection (LOD) and vehicle control (VC). Black column: 10 μM molnupiravir alone, grey columns: single compound treatments, striped columns: molnupiravir-based combinations. *p<0.05, **p<0.01, ***p<0.001, **** p<0.0001, representing significance over molnupiravir alone. Statistical significance between single agents and combinations are shown numerically. Data are represented as mean ± SD. Dashed lines, VC: vehicle control, LOD: limit of detection.

### Determination of IC_50_ of molnupiravir

Molnupiravir, which has since demonstrated clinical efficacy, was chosen as the basis compound to evaluate drug combinations as a treatment option against SARS-CoV-2 infection. Using pooled data from two testing sites, a dose response curve (Figure 3) of the fold change in genome copies following treatment with molnupiravir estimated IC_50_ at 11.1 μM (Figure 3). Similar IC_50_ of 9.2 μM was calculated for molnupiravir using infectious titer in apical wash (Figure 3c). Based on these data, the concentration of molnupiravir was set to 10 μM in combination treatments.

### Antiviral activity of molnupiravir-based combinations against SARS-CoV-2

Overall, 30 combinations were tested at various concentrations (48 different conditions, Table S2). Treatment of SARS-CoV-2 infected re-constituted epithelium with 10 μM molnupiravir alone resulted in a moderate 2 log reduction in viral RNA both in apical wash and intracellularly (Figure 3a and b, black column). The combination of molnupiravir at a submaximal dose with other compounds resulted in significantly higher antiviral activity measured by qPCR for apilimod, alisporivir, nafamostat, ivermectin, camostat and brequinar (Figure 3a, striped bars). These compounds showed limited antiviral effect alone (Figure 3a, grey bars) and the combined effect was also significantly better than treatment with molnupiravir alone. Interestingly, when looking at remaining infectious virus in apical wash, all combinations except molnupiravir and PF-00835231 showed a significant reduction compared to single treatments, many conditions resulting in no detectable infectious virus (Figure 3b). Combinations with ONO-3307, TO-195, and remdesivir, that did not exhibit a statistically significant benefit in the qPCR analysis, also presented with no detectable infectious virus at 72 hpi.

### Secretions of IP-10 and RANTES after combination treatment

The secretion of two major chemokines induced by SARS-CoV-2 infection, IP-10/CXCL-10 and RANTES/CXCL-5, was moderate, with an average of 4.5 and 9.2-fold increase over mock respectively. In the combination treatments, we did not observe any robust pattern of inhibition. IP-10 secretion was generally low, and only significantly reduced in combination treatment with remdesivir (p<0.0001). For nafamostat, ivermectin, ONO-3307, and brequinar no reduction was observed after the addition of molnupiravir (Figure S3a) while a slight non-significant reduction was observed for alisporivir and TO-195. A similar pattern was observed for RANTES (Figure S3b).

## Discussion

Despite current access to vaccination against SARS-CoV-2, safe and effective antiviral treatment options should be pursued and refined for human use. For optimal antiviral treatment, a drug combination should interfere with virus replication and disease progression at multiple crucial levels to provide a comprehensive treatment against both the virus infection and the resulting disease. Furthermore, risk of escape mutations that render virus-targeting drugs ineffective, is lower with a multidrug treatment approach, as observed during treatment against HCV and in antiretroviral treatment against HIV (36-38). In the current study, we have assessed the antiviral potential of compounds from the ReFrame and MMV drug libraries against SARS-CoV-2, both alone and in combination at two separate testing sites with a successful transfer of a standardized protocol in commercially available re-constituted airway epithelia, MucilAir™ (39). Importantly, this transposable approach facilitates both reproducible results and a clear distinction between antiviral and general toxic effect using the trans-epithelial electrical resistance (TEER) as a surrogate marker, an important feature of this organotypic testing system. Indeed, we observed increased toxicity beyond that of vehicle control after treatment with certain compounds only after infection with SARS-CoV-2. This observation points towards the potential of synergistic cellular toxicity between certain drugs and SARS-CoV-2 infection, highlighting the need for extensive toxicity testing prior and during to antiviral compound testing.

In the current study, the top five most active single treatments were all compounds that interfere directly with either viral proteases or the RdRp, i.e., PF-00835231, remdesivir, molnupiravir, nelfinavir, and ritonavir. Other protease inhibitors, ONO-3307 and TO-195 were also antiviral at the highest concentration tested (30 μM). PF-00835231 is the active metabolite of lufotrelvir, which is currently part of the pivotal ACTIV 3 clinical trial expected to report later in 2022. Other borderline antiviral compounds included digitoxin, narasin, nafamostat, and ivermectin with varying mechanisms of action (MoA). However, compounds with such borderline antiviral activity are likely ineffective clinically, as has been observed for ivermectin (40-42). Interestingly, antimalarial compounds, including the 4-aminoquinolines chloroquine, amodiaquine, and pyronaridine, did not show antiviral activity in this assay, in contrast with results from other cell lines. Combination treatment based on the orally bioactive RdRp inhibitor molnupiravir at 10 μM, resulted in varying level of viral inhibition with statistically significant improvement in antiviral activity for apilimod, alisporivir, nafamostat, ivermectin, camostat, and brequinar when evaluated by qPCR. Interestingly, all top 10 candidates, apart from PF-00835231, showed significant reduction in infectious titer in combination with molnupiravir compared to treatment with a single agent, highlighting the importance of comprehensive molecular and functional analysis of antiviral compounds. Here, we observed only moderate upregulation of the chemokines IP-10 and RANTES after infection with SARS-CoV-2 and a limited impact of combination treatments on their secretion. As a result, we were unable to discern any pattern of benefit.

In November 2021, Merck (MSD) and Ridgeback Biotherapeutics received emergency authorization from the American Food and Drug Administration for the use of molnupiravir against SARS-CoV-2 infection due to its excellent performance in clinical trials (43). Additionally, when combined with Favipiravir, antiviral activity of molnupiravir against SARS-CoV-2 in Syrian hamsters was enhanced (44), further highlighting the potential benefits of combination therapies. Molnupiravir is also orally bioactive, making its administration easy and straightforward. Furthermore, combination treatments could require lower dosages of single active compounds potentially providing additional therapeutic benefits.

In conclusion, the herein reported molnupiravir-based combination treatments increased antiviral activity against SARS-CoV-2 compared to single treatments in re-constituted nasal epithelium. Further studies focusing on the precise additive or synergistic effect of molnupiravir-based combinations are needed to provide a more comprehensive idea of their treatment potential.

## Author Contributions

Conceptualization, H.R.J., B.B., O.B.E. and S.C. Formal Analysis H.R.J., B.B., O.B.E. and S.C. Funding Acquisition T.W, S.C. Investigation, H.R.J., D.S., T.J., B.P., M.B., O.T., A.P. Methodology, H.R.J., B.B., O.B.E. and S.C. Project Administration M.R.-C., O.B.E., S.C. Resources K. S., T.W. Supervision M.R.-C., O.B.E., S.C. Validation H.R.J., B.B., O.B.E. and S.C Visualization, H.R.J. and B.B., Writing – Original Draft H.R.J., B.B., O.B.E. and S.C. Writing – Review & Editing, H.R.J., O.T., S.H., T.W., B.B., O.B.E., M.R.-C. and S.C.

## Acknowledgements

We thank Monalisa Chatterji, Robert Jordan and Betsy Russell from the Bill and Melinda Gates Foundation for advice and discussions. We thank Mitchell Hull and Kelli Kuhen from CALIBR for providing the ReFrame library and validation plates. This work was supported by funding to Medicines for Malaria Venture from the UK Foreign and Commonwealth Development Office, and the Bill and Melinda Gates Foundation (INV-016776).

## Figure Legends

**Figure S1:**
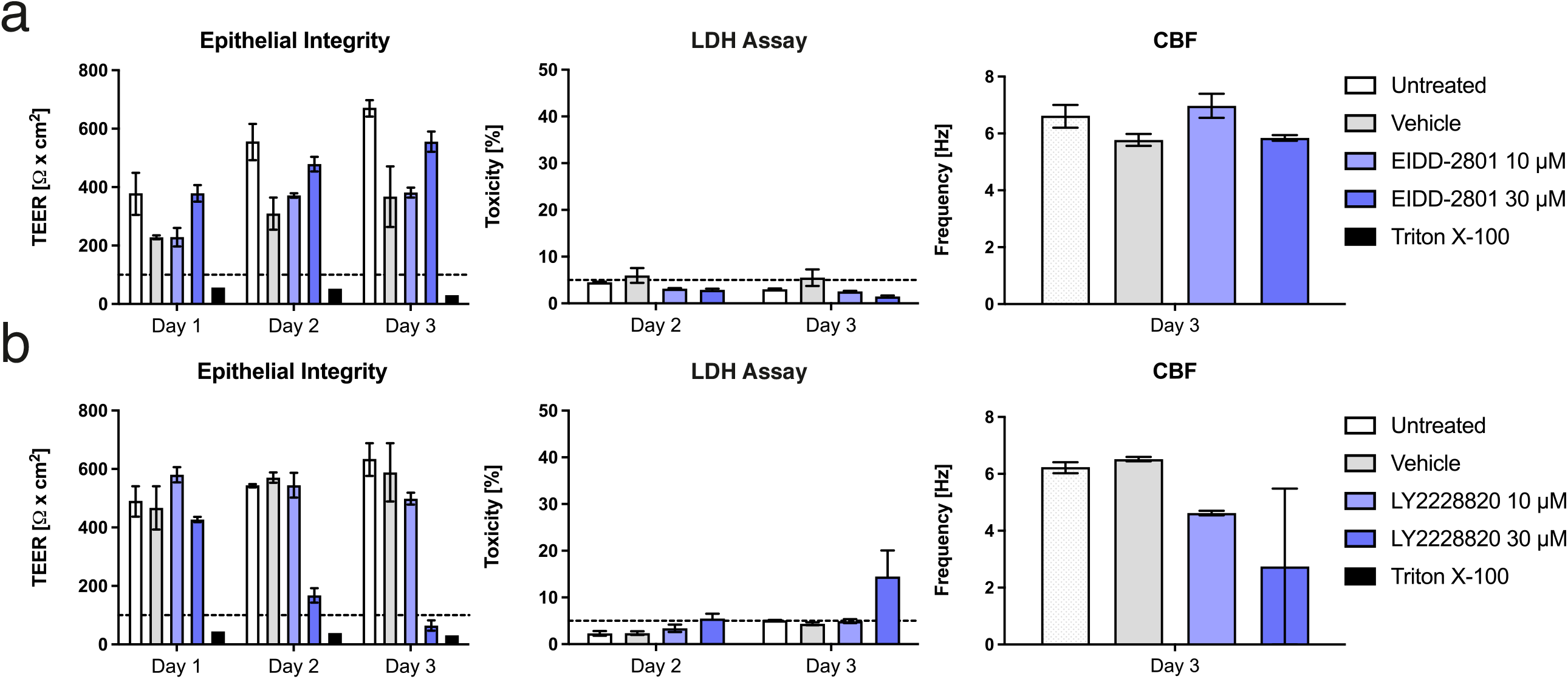
Determination of compound toxicity for dosage selection. Typical representation of parameters determining compound toxicity; epithelial integrity (TEER), LDH release, ciliary beating frequency (CBF) on days 1, 2 and 3, for **a)** molnupiravir (n=3) and **b)** LY2228820 (n=2). Molnupiravir is well tolerated at both 30 and 10 μM while LY2228820 exhibits toxicity at 30 μM: TEER is measured <100 Ohm.cm^2^ at day 3, indicating a loss of tissue integrity, cytotoxicity on day 3 >5 %, which corresponds to a physiological turnover of the nasal epithelial model MucilAir™ and CBF was strongly decreased. Data are represented as mean ± standard error of the mean (SEM).

**Figure S2:**
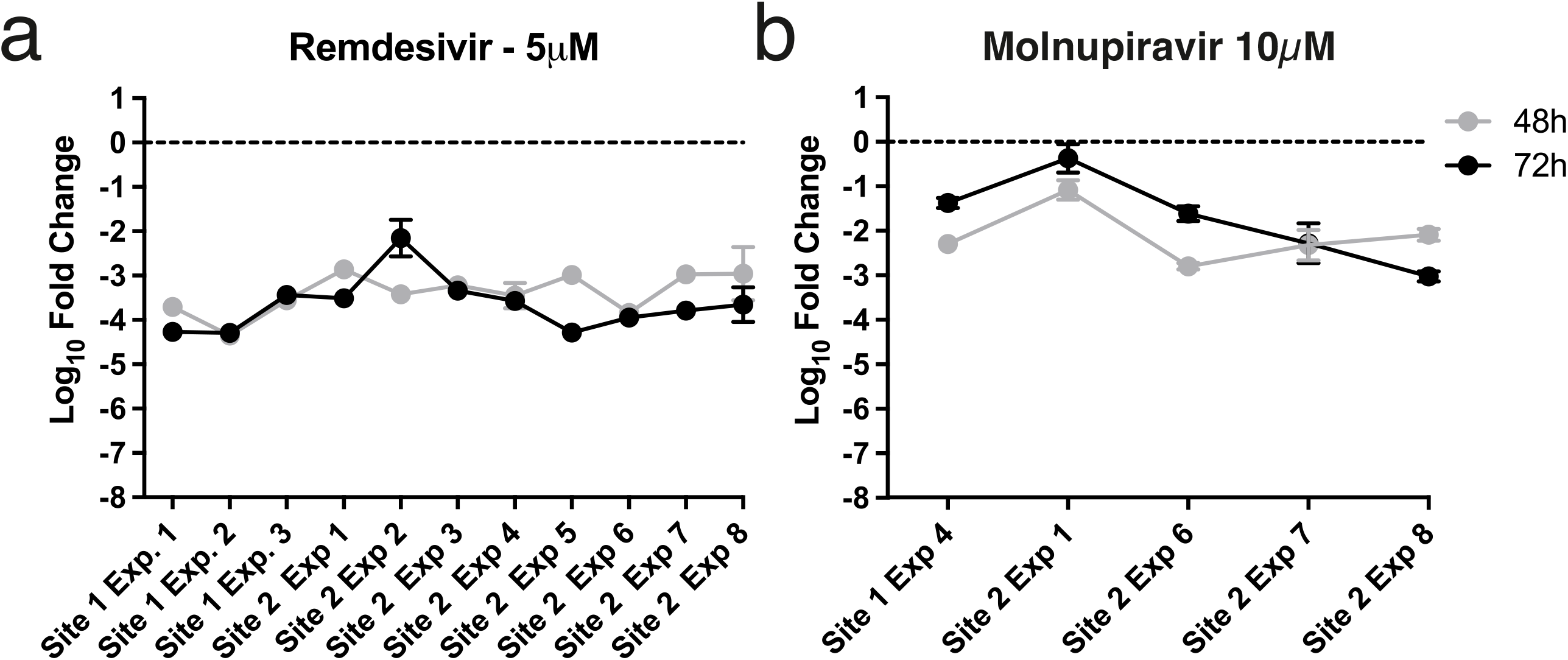
Reproducibility of antiviral compound testing in MucilAir™ nasal epithelia between two test sites. Repeated tests conducted with **a)** 5 μM remdesivir or **b)** 10 μM molnupiravir at two separate test locations using the same cell culture system and infection protocol. Data are represented as mean ± SEM, n=2-3.

**Figure S3:**
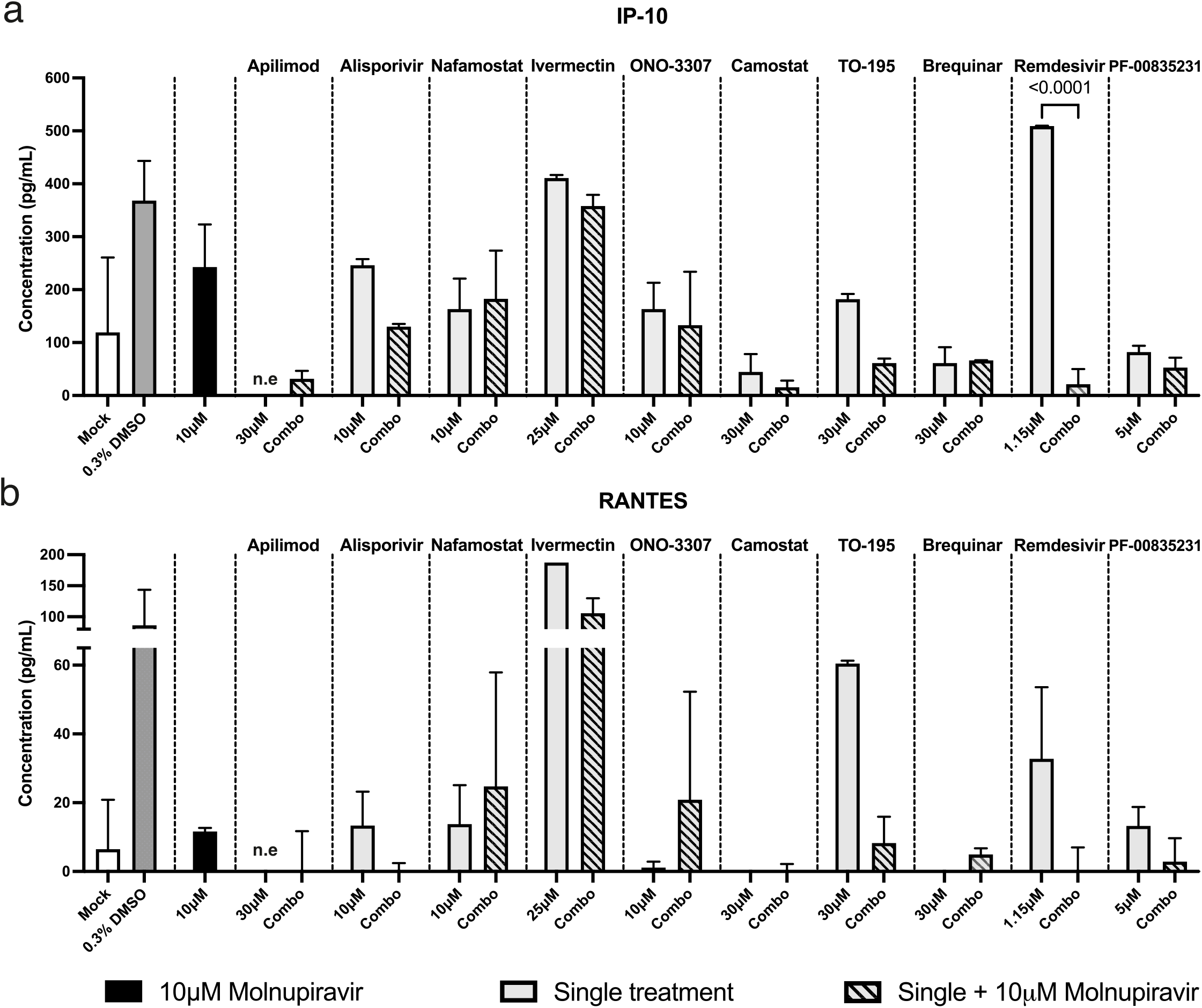
Secretion of IP-10 and RANTES after combination treatment. Secretion of the chemokines **a)** IP-10 (CXCL-10) and **b)** RANTES (CXCL-5) after molnupiravir-based combination treatment against SARS-CoV-2 as measured by ELISA (n=2-4). White column: Mock (uninfected control), dark grey column: infected vehicle control containing 0.3 % DMSO (VC), black column: 10 μM molnupiravir alone, light grey columns: single compound treatments, striped columns: molnupiravir-based combinations. *p<0.05, **p<0.01, ***p<0.001, **** p<0.0001, representing significance over molnupiravir alone. Statistical significance between single agents and combinations are represented numerically. Data are represented as mean ± SD. n.e: not evaluated.

