## Supplementary tables for "Molnupiravir combined with different repurposed drugs further inhibits SARS-CoV-2 infection in human nasal epithelium *in vitro*"

**Table S1: Single compounds tested on reconstituted nasal epithelium**. Maximum tolerated dose (≤30 µM) obtained during toxicity-testing and antiviral effects of different concentrations assessed at 48 and 72 hpi in apical wash and at 72 hpi intracellularly, data are represented as mean ± SD of $log_{10}$ reduction over vehicle control (VC). Toxic conditions, based on TEER< VC, have been excluded.

| Compound | MoA | MTD (µM) | Dosage (µM) | RT-qPCR - Nsp-14 ($Log_{10}$ reduction) | | | | | |
|---|---|---|---|---|---|---|---|---|---|
| | | | | **AW48h** | **±SD** | **AW72h** | **±SD** | **IC 72h** | **±SD** |
| 5-Aminolevulinic | Cancer detection | 30 | 30 | -0,633 | 0,345 | -0,380 | 0,126 | -2,163 | 0,017 |
| | | | 10 | -0,801 | 0,079 | -0,500 | 0,019 | -2,399 | 0,121 |
| Abacavir | RdRp inhibitor | 30 | 30 | -0,158 | 0,166 | -0,453 | 0,324 | -2,027 | 0,081 |
| | | | 10 | -0,092 | 0,677 | -0,568 | 0,034 | -2,241 | 0,026 |
| Albaconazole | Antifungal | 30 | 30 | 0,226 | 0,329 | -0,069 | 0,127 | -0,814 | 0,075 |
| | | | 10 | 0,376 | 0,309 | -0,088 | 0,104 | -0,723 | 0,060 |
| Alisporivir | Cyclophilin inhibitor | 10 | 10 | 1,404 | 0,362 | 0,856 | 0,180 | 1,099 | 0,309 |
| | | | 3 | -0,240 | 0,402 | -0,180 | 0,029 | 0,189 | 0,079 |
| Amiodarone | Potassium blocker | 30 | 30 | -0,352 | 0,071 | -0,308 | 0,133 | -0,202 | 0,522 |
| | | | 10 | -0,344 | 0,242 | -0,193 | 0,267 | 0,087 | 0,032 |
| Amodiaquine | Antimalarial | 10 | 10 | -0,024 | 0,125 | 0,047 | 0,057 | -0,137 | 0,017 |
| | | | 5 | 0.634 | 0.508 | 0.122 | 0.152 | -0.256 | 0.130 |
| | | | 1 | 0.241 | 0.253 | -0.174 | 0.147 | -0.273 | 0.030 |
| Apilimod | PIKfyve inhibitor | 30 | 30 | 1,280 | 0,326 | 0,394 | 0,184 | 0,849 | 0,109 |
| | | | 10 | 0,898 | 0,146 | -0,231 | 0,106 | 0,496 | 0,204 |
| | | | 0,36 | -0,041 | 0,138 | 0,041 | 0,022 | 0,645 | 0,045 |
| Apitolisib | mTOR,PI3K inhibitor | 1 | 0,1 | -0,076 | 0,206 | -0,090 | 0,119 | 0,134 | 0,034 |
| | | | 0,03 | -0,033 | 0,199 | -0,051 | 0,055 | 0,323 | 0,187 |
| Arbidol/Umifenovir | Antiviral | 30 | 30 | -0,073 | 0,029 | -0,373 | 0,317 | -0,979 | 0,577 |
| | | | 10 | -0,164 | 0,016 | -0,292 | 0,158 | -0,620 | 0,011 |
| Artesunate | Generation of free radicals | 25 | 25 | 0.693 | 0.394 | -0.268 | 0.360 | -0.130 | 0.180 |

| | | | | | | | | | |
|---|---|---|---|---|---|---|---|---|---|
| | | | 5 | 0.443 | 0.198 | 0.084 | 0.371 | -0.122 | 0.080 |
| | | | 1 | 0.338 | 0.269 | 0.014 | 0.260 | -0.009 | 0.093 |
| AT-511* | RdRp inhibitor | 30 | 30 | -0,260 | 0,390 | 0,583 | 0,026 | 1,222 | 0,487 |
| | | | 10 | 0,659 | 0,376 | 1,070 | 0,611 | 0,551 | 0,164 |
| AT-527 Isomer 1 | RdRp inhibitor | 30 | 30 | 1,1078 | 0,4524 | -1,8590 | 0,1602 | 0,812 | 0,038 |
| | | | 10 | 1,4575 | 0,3865 | -2,1661 | 0,1896 | 0,587 | 0,390 |
| | | | 3,3 | 0,1751 | 0,4422 | -1,9819 | 0,2365 | 0,533 | 0,278 |
| | | | 1,15 | 0,7606 | 0,5962 | -1,6468 | 0,5178 | 0,640 | 0,073 |
| | | | 0,36 | 0,7310 | 0,1814 | -2,0311 | 0,3835 | 0,591 | 0,176 |
| AT-527 Isomer 2 | RdRp inhibitor | 30 | 30 | 2,2226 | 0,4586 | -1,3302 | 0,3407 | 0,767 | 0,232 |
| | | | 10 | 1,5709 | 0,3411 | -1,7732 | 0,3071 | 0,311 | 0,422 |
| | | | 3,3 | 1,2663 | 0,3777 | -1,9749 | 0,0989 | -0,073 | 0,152 |
| | | | 1,15 | 1,0611 | 0,2845 | -2,1314 | 0,1110 | -0,057 | 0,257 |
| | | | 0,36 | 0,8906 | 0,6008 | -1,6975 | 0,6291 | -0,211 | 0,434 |
| AZD-8055 | mTOR1,2 inhibitor | 3,3 | 3 | 0,758 | 0,019 | -1,669 | 0,083 | -0,340 | 0,038 |
| | | | 1 | 1,060 | 0,545 | -1,542 | 0,006 | -0,244 | 0,230 |
| Azithromycin | Bacterial protein synthesis inhibitor | 25 | 25 | 0.735 | 0.707 | 0.149 | 0.564 | 0.371 | 0.406 |
| | | | 5 | 0.078 | 0.321 | -0.277 | 0.457 | -0.117 | 0.069 |
| | | | 1 | 0.211 | 0.176 | -0.285 | 0.039 | -0.050 | 0.091 |
| Baricitinib | JAK inhibitor | 30 | 30 | -0,071 | 0,156 | -0,260 | 0,032 | -0,531 | 0,328 |
| | | | 10 | -0,109 | 0,172 | -0,199 | 0,039 | -0,301 | 0,075 |
| Bemcentinib/ R428 | AXL inhibitor | 30 | 30 | -0,147 | 0,036 | -0,912 | 0,304 | -2,340 | 0,085 |
| | | | 10 | -0,095 | 0,392 | -0,499 | 0,021 | -1,332 | 1,149 |
| Brequinar | DHODH inhibitor | 30 | 30 | 0,417 | 0,375 | 0,253 | 0,355 | 0,066 | 1,897 |
| | | | 10 | -0,420 | 0,221 | -0,176 | 0,085 | -1,665 | 0,232 |
| Bromhexine | Mucolytic drug | 30 | 30 | -0,361 | 0,168 | -0,720 | 0,147 | -2,086 | 0,138 |
| | | | 10 | -0,823 | 0,064 | -0,777 | 0,381 | -2,059 | 0,075 |
| Camostat mesylate | Serine protease inhibitor | 30 | 30 | 1,352 | 0,227 | 0,767 | 0,210 | 0,998 | 0,223 |
| | | | 10 | 0,747 | 0,782 | 0,542 | 0,045 | 0,690 | 0,183 |

| | | | | | | | | | |
|---|---|---|---|---|---|---|---|---|---|
| | | | 3,3 | -0,250 | 0,198 | 0,253 | 0,382 | 0,474 | 0,154 |
| | | | 1,15 | -0,314 | 0,135 | 0,155 | 0,457 | 0,434 | 0,066 |
| | | | 0,36 | -0,095 | 0,148 | 0,138 | 0,425 | 0,342 | 0,057 |
| Cepharantine | Anti-inflammatory | 10 | 1,15 | -0,372 | 0,067 | -0,256 | 0,060 | 0,455 | 0,048 |
| | | | 0,1 | 0,112 | 0,241 | 0,069 | 0,273 | 0,168 | 0,369 |
| Chloroquine | Antimalarial | 5 | 5 | 0.448 | 0.218 | 0.066 | 0.403 | -0.268 | 0.145 |
| | | | 1 | 0.442 | 0.054 | -0.166 | 0.080 | -0.322 | 0.060 |
| Daclatasvir | HCV Antiviral | 10 | 10 | 0.484 | 1.192 | -0.037 | 0.054 | 0.123 | 0.104 |
| | | | 5 | 0.467 | 0.516 | -0.097 | 0.087 | -0.142 | 0.260 |
| | | | 3 | 0,326 | - | -1,726 | 0,211 | -0,296 | 0,036 |
| | | | 1 | 0,169 | 0,328 | -1,414 | 0,196 | -0,533 | 0,064 |
| Dacomitinib | EGFR inhibitor | 10 | 10 | -0,308 | 0,064 | -0,551 | 0,049 | -2,240 | 0,023 |
| Dalbavancin | Antibiotic | 30 | 60 | 0,142 | 0,557 | -0,223 | 0,460 | 0,044 | 0,136 |
| | | | 20 | 0,186 | 0,095 | -0,184 | 0,075 | -0,190 | 0,087 |
| Dalcetrapib | CETP inhibitor | 30 | 30 | -0,566 | 0,032 | -0,526 | 0,179 | -2,082 | 0,132 |
| | | | 10 | -0,811 | 0,204 | -0,491 | 0,083 | -2,339 | 0,057 |
| Dexamethasone | Glucocorticoid | 30 | 30 | -0,166 | 0,019 | 0,043 | 0,016 | 2,296 | 0,028 |
| | | | 10 | -0,135 | 0,079 | 0,086 | 0,026 | -0,127 | 0,011 |
| Digitoxin | Cardiac glycoside | 3,3 | 0,36 | 3,303 | 0,632 | 2,540 | 0,913 | 3,805 | 1,760 |
| | | | 0,1 | 0,167 | 0,149 | 0,077 | 0,045 | 0,424 | 0,027 |
| | | | 0,003 | -0,060 | 0,293 | -0,056 | 0,137 | 0,136 | 0,182 |
| Digoxin | Na-K ATPase inhibitor | 0,1 | 0,1 | 0,286 | 0,030 | -0,509 | 0,062 | -0,297 | 0,132 |
| | | | 0,03 | 0,259 | 0,037 | -0,185 | 0,019 | 0,135 | 0,300 |
| Ebastine | Antihistamine | 30 | 30 | -0,299 | 0,083 | -1,544 | 0,528 | -0,238 | 0,166 |
| | | | 10 | -0,245 | 0,079 | -1,134 | 0,192 | -0,126 | 0,081 |
| Ebselen | Glutathione peroxidase mimic | 30 | 60[a] | 0,135 | 0,181 | -0,159 | 0,027 | 0,140 | 0,166 |
| | | | 20 | -0,402 | 0,036 | -0,381 | 0,181 | -0,166 | 0,117 |
| EIDD-2801 | RdRp inhibitor | 30 | 30 | 3,197 | 0,859 | 3,352 | 0,237 | 4,891 | 0,303 |
| | | | 20 | 3,219 | 0,071 | 3,494 | 0,439 | 3,551 | 1,373 |

| | | | | | | | | | |
|---|---|---|---|---|---|---|---|---|---|
| | | | 10 | 2,124 | 0,627 | 1,751 | 0,999 | 1,983 | 0,855 |
| | | | 5 | 1,376 | 0,004 | 1,157 | 0,165 | 1,072 | 0,331 |
| | | | 3,3 | 0,391 | 0,817 | 0,388 | 0,197 | 0,870 | 0,024 |
| | | | 2,5 | 0,755 | 0,017 | 0,796 | 0,068 | 0,988 | 0,307 |
| | | | 1,15 | -0,178 | 0,794 | 0,296 | 0,190 | 0,731 | 0,182 |
| | | | 0,36 | -0,127 | 0,676 | -0,131 | 0,342 | 0,410 | 0,129 |
| Emetine | Anti-protozoal/emetic | 1 | 1 | 1,704 | 0,471 | 1,234 | 0,312 | 1,330 | 0,321 |
| | | | 0,3 | 0,973 | 0,154 | 0,256 | 0,328 | 0,348 | 0,192 |
| Ezlopitant | NK1 receptor antagonist | 10 | 10 | 0,095 | 0,558 | 0,069 | 0,145 | 0,503 | 0,084 |
| | | | 3,3 | 0,277 | 0,128 | 0,210 | 0,100 | 0,465 | 0,126 |
| | | | 1,15 | -0,073 | 0,099 | -0,204 | 0,024 | 0,369 | 0,061 |
| | | | 0,36 | -0,184 | 0,100 | -0,164 | 0,103 | 0,439 | 0,054 |
| Favipiravir | RdRp inhibitor | 1 | 1 | 0.362 | 0.570 | 0.068 | 0.075 | 0.316 | 0.136 |
| Fenofibrate | Activates PPARα | 30 | 30 | -0,597 | 0,017 | -0,514 | 0,170 | -2,205 | 0,000 |
| | | | 10 | -0,355 | 0,368 | -0,422 | 0,019 | -2,175 | 0,192 |
| Ferroquine | Antimalarial | 5 | 5 | -0.382 | 0.360 | -0.361 | 0.042 | -0.251 | 0.228 |
| | | | 1 | -0.531 | 0.021 | -0.110 | 0.182 | -0.090 | 0.231 |
| Fingolimod | Sphingosine 1-phosphate receptor modulator | 30 | 30 | -0,028 | 0,115 | -0,548 | 0,040 | -2,234 | 0,019 |
| | | | 10 | -0,269 | 0,064 | -0,648 | 0,355 | -1,979 | 0,089 |
| Fluvoxamine | SSRI | 30 | 30 | 1,206 | 0,613 | 0,013 | 0,098 | -0,363 | 1,780 |
| | | | 10 | -0,036 | 0,275 | -0,366 | 0,038 | -2,002 | 0,211 |
| Halofantrine | Antimalarial | 5 | 5 | -0.063 | 0.787 | 0.203 | 0.016 | 0.206 | 0.172 |
| Hydroxychloroquine | Metabolite of chloroquine | 25 | 25 | 0.050 | 0.092 | -0.213 | 0.051 | 0.020 | 0.159 |
| | | | 5 | 0.237 | 0.434 | -0.039 | 0.301 | -0.251 | 0.228 |
| | | | 1 | 0.728 | 0.716 | -0.142 | 0.182 | -0.090 | 0.231 |
| Homoharringtonine | Protein synthesis | 3,3 | 0,1 | 3,410 | 0,147 | 1,158 | 0,206 | 3,707 | 0,245 |
| | | | 0,03 | 0,923 | 0,032 | -1,520 | 0,123 | 0,259 | 0,285 |
| Imatinib | Tyrosine kinase inhibitor | 5 | 5 | 0.461 | 0.560 | -0.161 | 0.336 | 0.133 | 0.259 |
| | | | 1 | 0.201 | 0.332 | -0.374 | 0.289 | 0.003 | 0.116 |

| | | | | | | | | | |
|---|---|---|---|---|---|---|---|---|---|
| Indomethacin | NSAID | 25 | 25 | 0.158 | 0.411 | 0.027 | 0.176 | 0.263 | 0.068 |
| Interferon beta | Interferon | 100 IU/ml | 200 | 1,233 | 0,921 | 1,231 | 0,232 | 0,608 | 0,349 |
| | | | 100 | 0,764 | 0,334 | -0,290 | 0,126 | 0,851 | 0,019 |
| | | | 75 | 0,630 | 0,630 | -0,200 | 0,189 | 0,747 | 0,251 |
| Interferon lambda | Interferon | 50 IU/ml | 50 | 0,449 | 0,038 | -0,787 | 0,066 | 0,422 | 0,221 |
| | | | 10 | 0,559 | 0,138 | -1,122 | 0,732 | 0,051 | 0,115 |
| IPI-549 | PI3K inhibitor | 3 | 3 | 0,619 | 0,302 | -0,321 | 0,051 | -2,678 | 0,055 |
| | | | 1 | 0,072 | 0,313 | -0,485 | 0,023 | -2,408 | 0,330 |
| Irinotecan | Cytotoxic alkaloid | 10 | 10 | 0,428 | 0,557 | 0,098 | 0,296 | -0,015 | 0,189 |
| | | | 3 | 0,096 | 0,247 | -0,194 | 0,096 | -0,346 | 0,062 |
| Ivermectin | Anta-parasitic | 25 | 25 | 0.841 | 0.910 | 1.253 | 0.242 | 1.213 | 0.040 |
| | | | 10 | 0,851 | 0,153 | 1,303 | 0,184 | 1,104 | 0,262 |
| | | | 5 | 1.139 | 0.321 | 0.223 | 0.228 | 0.260 | 0.067 |
| | | | 1 | 1.257 | 0.221 | 0.102 | 0.202 | -0.183 | 0.135 |
| Leflunomide | Pyrimidine synthesis inhibitor | 30 | 3,3 | -0,004 | 0,260 | -0,087 | 0,103 | 0,361 | 0,295 |
| | | | 1,15 | 0,031 | 0,246 | 0,312 | 0,448 | 0,176 | 0,424 |
| | | | 0,36 | -0,199 | 0,253 | 0,069 | 0,069 | 0,360 | 0,422 |
| Lopinavir | Protease inhibitor | 3,3 | 3,3 | 0,147 | 0,330 | 0,038 | 0,077 | 0,416 | 0,191 |
| Losartan | Angiotensin II receptor antagonists | 30 | 30 | -0,244 | 0,295 | 0,157 | 0,168 | 0,033 | 0,108 |
| | | | 10 | 0,434 | 0,289 | -0,013 | 0,226 | 0,130 | 0,148 |
| | | | 3,3 | -0,052 | 0,183 | -0,141 | 0,179 | -0,063 | 0,071 |
| | | | 1,15 | -0,231 | 0,650 | -0,129 | 0,066 | -0,018 | 0,117 |
| | | | 0,36 | -0,185 | 0,156 | -0,038 | 0,215 | 0,035 | 0,046 |
| Lumefantrine | Antimalarial | 25 | 25 | 0.075 | 0.232 | 0.380 | 0.140 | 0.056 | 0.070 |
| | | | 5 | 0.797 | 0.913 | -0.247 | 0.098 | -0.010 | 0.092 |
| | | | 1 | 0.356 | 0.235 | -0.246 | 0.188 | 0.138 | 0.088 |
| LY2228820 | Protein kinase inhibitor | 10 | 10 | 0,372 | 0,430 | 0,155 | 0,045 | 0,123 | 0,126 |
| | | | 3 | 0,167 | 0,206 | -0,049 | 0,122 | -0,008 | 0,187 |
| Manidipine | Calcium Channel blocker | 30 | 30 | -0,181 | 0,164 | -0,115 | 0,029 | -0,592 | 0,898 |

| | | | | | | | | | |
|---|---|---|---|---|---|---|---|---|---|
| | | | 10 | -0,305 | 0,146 | -0,045 | 0,143 | 0,433 | 0,283 |
| (+)Mefloquine | Antimalarial | 25 | 25 | -0.101 | 0.241 | -0.065 | 0.296 | -0.241 | 0.067 |
| | | | 5 | 0.251 | 0.219 | -0.084 | 0.269 | -0.058 | 0.203 |
| | | | 1 | 1.059 | 0.343 | 1.218 | 0.177 | 0.892 | 0.139 |
| Midostaurine / PKC412 | Kinase inhibitor | 1 | 0,3 | -0,231 | 0,222 | 0,033 | 0,100 | -0,130 | 0,002 |
| Mitoguazone | Polyamine synthesis | 10 | 10 | 1,055 | 1,671 | 0,580 | 0,020 | 1,436 | 0,581 |
| | | | 3 | 1,771 | 0,414 | 0,043 | 0,248 | 0,650 | 0,138 |
| Mycophenolic Acid | Immunosuppressant | 30 | 30 | -0,079 | 0,264 | 0,008 | 0,026 | 0,385 | 0,530 |
| | | | 10 | -0,268 | 0,358 | -0,099 | 0,083 | 0,188 | 0,298 |
| N-desethylamodiaquine | Metabolite of amodiaquine | 1 | 1 | 0.755 | 0.320 | 0.206 | 0.302 | 0.074 | 0.055 |
| Nafamostat | Serine protease inhibitor | 30 | 30 | 3,354 | - | 1,621 | 0,722 | 4,556 | 0,032 |
| | | | 10 | 2,366 | 0,751 | 0,619 | 2,122 | 1,220 | 2,125 |
| Nanchangmycin | Antibiotic | 3,3 | 1 | 1,804 | 0,272 | -0,344 | 0,275 | 0,571 | 0,292 |
| | | | 0,3 | 1,117 | 0,298 | -0,328 | 0,021 | 0,190 | 0,255 |
| Narasin | Coccidiostat/antibacterial | 1 | 1 | 2,985 | 0,376 | 1,489 | 0,811 | 0,519 | 0,047 |
| | | | 0,3 | 2,188 | - | 1,197 | - | 0,053 | - |
| Nelfinavir | Protease inhibitor | 30 | 30 | 3,259 | 0,344 | 3,179 | 0,162 | 5,154 | 0,903 |
| | | | 10 | 2,408 | 0,650 | 2,404 | 0,588 | 1,440 | 0,593 |
| | | | 3,3 | 0,323 | 0,406 | 0,277 | 0,220 | 0,552 | 0,045 |
| | | | 1,15 | -0,054 | 0,008 | 0,099 | 0,317 | 0,457 | 0,038 |
| | | | 0,36 | -0,040 | 0,265 | -0,155 | 0,147 | 0,491 | 0,042 |
| Nevibolol hydrochloride | Beta 1 receptor antagonist | 10 | 10 | -0,053 | 0,171 | 0,104 | 0,172 | 0,288 | 0,182 |
| | | | 3,3 | 0,263 | 0,180 | -0,068 | 0,123 | 0,194 | 0,102 |
| | | | 1,15 | -0,259 | 0,181 | -0,145 | 0,012 | 0,089 | 0,080 |
| | | | 0,36 | 0,007 | 0,171 | -0,244 | 0,109 | 0,191 | 0,120 |
| | | | 0,1 | 0,013 | 0,254 | 0,053 | 0,173 | 0,280 | 0,107 |
| Niclosamide | Antiparasitic | <25 | 10 | -0,561 | 0,071 | 0,124 | 0,018 | 0,936 | 0,181 |
| | | | 5 | -0.213 | 0.226 | 0.546 | 0.158 | 0.331 | 0.067 |
| | | | 3 | -0,154 | 0,083 | -1,300 | 0,009 | -0,092 | 0,019 |

| | | | | | | | | | |
|---|---|---|---|---|---|---|---|---|---|
| | | | 1 | -0.286 | 0.335 | 0.124 | 0.201 | 0.184 | 0.028 |
| | | | 1 | 0,258 | 0,168 | -1,663 | 0,381 | -0,247 | 0,179 |
| Nifedipine | Calcium channel blocker | 30 | 30 | 0,233 | 0,144 | 0,147 | 0,195 | 0,698 | 0,056 |
| | | | 10 | 0,477 | 0,330 | 0,192 | 0,251 | 0,199 | 0,288 |
| | | | 3,3 | 0,441 | 0,353 | 0,179 | 0,117 | 0,316 | 0,286 |
| | | | 1,15 | 0,325 | 0,490 | -0,045 | 0,236 | 0,130 | 0,292 |
| | | | 0,36 | -0,058 | 0,175 | -0,011 | 0,090 | 0,122 | 0,418 |
| Nitazoxanide | Antiparasitic | 10 | 10 | -0.350 | 0.154 | 0.526 | 0.371 | 0.197 | 0.095 |
| | | | 5 | 0.152 | 0.406 | 0.065 | 0.097 | 0.039 | 0.047 |
| | | | 1 | 0.043 | 0.285 | -0.097 | 0.104 | 0.011 | 0.106 |
| ONO-3307 | Protease | 30 | 30 | 3,459 | 0,016 | 2,491 | 0,741 | 2,771 | 0,494 |
| | | | 10 | 1,896 | 0,547 | 0,579 | 0,282 | 0,695 | 0,590 |
| ONO-5334 | Protease inhibitor | 30 | 30 | 0,192 | 0,104 | 0,170 | 0,021 | -0,589 | 0,057 |
| | | | 10 | 0,026 | 0,148 | 0,298 | 0,229 | -0,136 | 0,119 |
| Ozanimod | S1P receptor modulator | | 10 | 0,002 | 0,096 | 0,355 | 0,185 | -0,274 | 0,081 |
| Patamostat mesylate | Protease inhibitor | 30 | 30 | 1,180 | 1,377 | 0,452 | 0,862 | 0,789 | 0,573 |
| | | | 10 | 1,488 | 0,459 | 0,602 | 0,166 | 0,473 | 0,373 |
| PF-00835231 | Protease inhibitor | 30 | 30 | 3,418 | 0,216 | 4,223 | 0,332 | 5,063 | 0,106 |
| | | | 10 | 3,674 | 0,518 | 4,334 | 0,022 | 5,072 | 0,068 |
| | | | 5 | 2,754 | 0,232 | 3,665 | 0,496 | 4,126 | 0,051 |
| Piperaquine | Antimalarial | 25 | 25 | 0.195 | 0.268 | -0.091 | 0.228 | 0.002 | 0.135 |
| Pyronaridine | Antimalarial | 5 | 5 | -0.561 | 0.036 | -0.290 | 0.062 | -0.268 | 0.145 |
| | | | 1 | 0.194 | 0.140 | -0.119 | 0.152 | -0.322 | 0.060 |
| Quinacrine | Antiplasmodial | 5 | 5 | 0.053 | 0.273 | -0.322 | 0.270 | 0.066 | 0.084 |
| | | | 1 | 0.286 | 0.239 | 0.309 | 0.420 | -0.013 | 0.211 |
| Quinine | Antimalarial | 25 | 25 | 0.710 | 0.315 | -0.158 | 0.206 | -0.064 | 0.109 |
| | | | 5 | 0.413 | 0.142 | 0.307 | 0.347 | -0.028 | 0.086 |
| | | | 1 | 0.177 | 0.287 | -0.197 | 0.327 | -0.135 | 0.038 |
| Remdesivir | RdRp inhibitor | 10 | 10 | 3,326 | 0,723 | 2,360 | 0,726 | 5,872 | 0,028 |

| | | | | | | | | | |
|---|---|---|---|---|---|---|---|---|---|
| | | | 5 | 3,054 | 0,986 | 3,188 | 0,739 | 5,458 | 1,083 |
| | | | 3,3 | 3,226 | 1,050 | 3,056 | 0,105 | 5,911 | 0,725 |
| | | | 1,15 | 3,076 | 0,804 | 2,599 | 0,813 | 4,339 | 2,300 |
| | | | 0,36 | 2,129 | 0,836 | 1,661 | 0,273 | 2,546 | 1,262 |
| Risdiplam | SMA treatment | 10 | 10 | -0,311 | 0,199 | -0,023 | 0,137 | 0,173 | 0,060 |
| | | | 3 | -0,032 | 0,111 | -0,275 | 0,053 | 0,042 | 0,006 |
| Ritonavir | HIV protease inhibitor | 30 | 30 | 2,296 | 0,339 | 2,336 | 0,588 | 2,170 | 0,767 |
| | | | 10 | 1,498 | 0,306 | 0,456 | 0,241 | 0,514 | 0,047 |
| | | | 0,36 | 0,047 | 0,205 | 0,059 | 0,142 | 0,152 | 0,097 |
| Salinomycin | Coccidiostat/antibacterial | 3,3 | 1 | 0,101 | 0,275 | -0,833 | 0,502 | 0,100 | 0,034 |
| | | | 0,3 | 0,095 | 0,147 | -1,195 | 0,072 | 0,095 | 0,300 |
| Samotolisib | PI3K, mTOR inhibitor | 3 | 3 | 0,726 | 0,385 | 0,088 | 0,145 | -0,971 | 1,205 |
| | | | 1 | 0,279 | 0,613 | -0,053 | 0,119 | -0,789 | 0,236 |
| Silmitasertib/CX-4945 | Kinase inhibitor, protein kinase CK2 | 3 | 3 | -0,188 | 0,102 | -0,497 | 0,040 | -2,586 | 0,126 |
| | | | 1 | -0,758 | 0,253 | -0,517 | 0,060 | -2,923 | 0,709 |
| Sofosbuvir | HCV RdRp inhibitor | 30 | 30 | -0,174 | 0,119 | -1,143 | 0,230 | -0,192 | 0,140 |
| | | | 25 | 0.080 | 0.316 | 0.029 | 0.097 | 0.134 | 0.084 |
| | | | 10 | -0,389 | 0,134 | -1,216 | 0,383 | -0,013 | 0,070 |
| | | | 5 | 0.263 | 0.214 | 0.031 | 0.085 | 0.253 | 0.081 |
| Sorafenib | Raf/Mek/Erk | 10 | 10 | 0,001 | 0,036 | 0,139 | 0,217 | 0,647 | 0,053 |
| | | | 3 | 0,012 | 0,017 | 0,083 | 0,062 | 0,462 | 0,400 |
| Tenofovir | RdRp inhibitor | 30 | 30 | 0,042 | 0,363 | 0,138 | 0,154 | 0,196 | 0,139 |
| Tetrandrine | Calcium Channel blocker | 10 | 10 | 0,061 | 0,230 | -1,503 | 0,155 | 0,173 | 0,462 |
| | | | 3 | -0,069 | 0,264 | -1,700 | 0,192 | -0,432 | 0,160 |
| | | | 1 | 0.433 | 0.247 | -0.036 | 0.056 | 0.227 | 0.043 |
| Tizoxanide | Metabolite of nitazoxanide | 25 | 25 | 0.005 | 0.430 | 0.330 | 0.139 | 0.220 | 0.064 |
| TO-195 | Protease inhibitor | 30 | 30 | 2,619 | 0,302 | 2,006 | 0,640 | 2,404 | 0,851 |
| | | | 10 | 0,531 | 0,302 | -0,038 | 0,325 | 0,445 | 0,390 |
| Toremifene | | 30 | 10 | -0,058 | 0,137 | 0,017 | 0,208 | 0,405 | 0,104 |

| | | | | | | | | | |
|---|---|---|---|---|---|---|---|---|---|
| | Estrogen receptor modulator (SERM) | | 0,36 | -0,073 | 0,557 | 0,121 | 0,131 | 0,260 | 0,025 |
| Trimethoprim | Bacterial DHFR inhibitor | 25 | 25 | 0.010 | 0.714 | 0.334 | 0.161 | 0.316 | 0.097 |
| VBY-825 | Protease inhibitor | 30 | 30 | 0,680 | 0,999 | 0,214 | 0,210 | -0,345 | 0,215 |
| | | | 10 | 0,440 | 0,030 | 0,047 | 0,159 | -0,569 | 0,038 |
| Verapamil | Calcium Channel blocker | 30 | 30 | 0,404 | 0,430 | -0,782 | 0,221 | 0,494 | 0,609 |
| | | | 25 | 0.712 | 0.120 | 0.021 | 0.062 | 0.217 | 0.040 |
| | | | 10 | -0,025 | 0,040 | -1,053 | 0,481 | 0,089 | 0,015 |
| | | | 5 | 0.207 | 0.158 | -0.085 | 0.057 | -0.082 | 0.351 |
| Vidofludimus | DHODH inhibitor | 10 | 10 | -0,6560 | 0,6380 | -0,2425 | 0,3414 | 0,052 | 0,336 |
| | | | 5 | -0.389 | 0.612 | 0.086 | 0.063 | 0.063 | 0.143 |
| | | | 3,3 | -0,1909 | 0,2243 | -0,5450 | 0,1613 | 0,497 | 0,328 |
| | | | 1,15 | -0,6736 | 0,4572 | -0,0157 | 0,6856 | 0,020 | 0,175 |
| | | | 0,36 | -0,6866 | 0,5035 | 0,2979 | 0,5076 | 0,227 | 0,095 |
| | | | 0,1 | -0,9104 | 0,2870 | -0,1446 | 0,3116 | 0,250 | 0,100 |
| Z LVG CHN2 | Cysteine proteinase inhibitor | 30 | 10 | -0,295 | 0,148 | -0,166 | 0,116 | -0,464 | 0,121 |

MoA: Mechanism of Action, AW: Apical Wash, IC: Intracellular, SD: Standard Deviation
*tested according to the protocol of https://www.biorxiv.org/content/10.1101/2020.08.11.242834v2

**Table S2: Molnupiravir-based combinations tested on reconstituted nasal epithelium**. Antiviral effects of different concentrations assessed at 48 and 72 hpi in apical wash and at 72 hpi intracellularly, data are represented as mean ± SD of $log_{10}$ reduction over vehicle control (VC). Toxic conditions, based on TEER< VC, have been excluded.

| Compound 1 | Compound 2 | MoA (1/2) | Conc.1 (µM) | Conc. 2 (µM) | AW 48h | (±)SD | AW 72h | (±)SD | IC 72h | (±)SD |
|---|---|---|---|---|---|---|---|---|---|---|
| Molnupiravir | Alisporivir | RdRp/Cyclophilin | 10 | 10 | 2,728 | 0,486 | 3,732 | 0,286 | 4,292 | 0,110 |
| Molnupiravir | Amodiaquine | RdRp/Antimalarial | 10 | 10 | 2,407 | 0,154 | 2,343 | 0,587 | 3,034 | 0,107 |
| Molnupiravir | Apilimod | RdRp/PIKfyve | 10 | 30 | 2,613 | 0,032 | 3,560 | 0,431 | 4,517 | 0,419 |
| Molnupiravir | Baricitinib | RdRp/JAK | 10 | 30 | 1,926 | 0,206 | 1,327 | 0,051 | 1,734 | 0,769 |
| | | | 10 | 10 | 1,379 | 0,223 | 0,940 | 0,112 | 1,022 | 0,054 |
| Molnupiravir | Brequinar | RdRp/DHODH | 10 | 30 | 2,278 | 0,123 | 2,986 | 0,339 | 3,609 | 0,161 |
| Molnupiravir | Camostat | RdRp/TMPRSS2 | 30 | 30 | 3,760 | 0,759 | 3,848 | 0,862 | 5,431 | 0,140 |
| | | | 10 | 10 | 3,051 | 0,119 | 3,437 | 0,124 | 3,439 | 0,465 |
| Molnupiravir | Dexamethasone | RdRp/Glucocorticoid | 10 | 30 | 2,018 | 0,564 | 2,059 | 0,336 | 2,296 | 0,028 |
| Molnupiravir | Ebselen | RdRp/Antioxidant | 10 | 30 | 1,859 | 0,085 | 2,303 | 0,077 | 3,208 | 0,526 |
| Molnupiravir | Emetine | RdRp/Antiprotozoal | 10 | 1 | 1,844 | 0,898 | 2,783 | 1,056 | 3,886 | 0,858 |
| Molnupiravir | Fluvoxamine | RdRp/SSRI | 10 | 30 | 2,674 | 0,233 | 3,011 | 0,658 | 3,111 | 0,402 |
| Molnupiravir | Homoharringtonine | RdRp/Anti-tumor | 10 | 0,1 | 3,538 | 0,247 | 3,395 | 0,080 | 4,950 | 0,192 |
| Molnupiravir | IFN beta (IU/ml) | RdRp/Interferon | 30 | 200 | 3,132 | 0,374 | 3,966 | 0,421 | 4,145 | 0,839 |

| | | | | | | | | | | |
|---|---|---|---|---|---|---|---|---|---|---|
| | | | 10 | 200 | 2,575 | 0,419 | 2,874 | 0,116 | 3,066 | 0,535 |
| | | | 10 | 100 | 2,621 | 0,140 | 2,878 | 0,318 | 2,634 | 0,736 |
| Molnupiravir | Ivermectin | RdRp/Antiparasitic | 30 | 30 | 3,225 | 0,143 | 3,885 | 0,119 | 5,349 | 0,272 |
| | | | 10 | 25 | 2,618 | 0,051 | 4,236 | 0,063 | 4,882 | 0,321 |
| | | | 10 | 10 | 3,031 | 0,366 | 2,783 | 0,346 | 3,096 | 0,164 |
| Molnupiravir | Mefloquine | RdRp/Antimalarial | 10 | 25 | 2,724 | 0,296 | 2,812 | 0,216 | 4,114 | 0,929 |
| Molnupiravir | Nafamostat | RdRp/TMPRSS2 | 30 | 30 | 3,891 | 0,346 | 3,420 | 0,804 | 5,718 | 0,773 |
| | | | 10 | 30 | 2,979 | 0,230 | 4,284 | 0,270 | 4,542 | 0,151 |
| | | | 10 | 10 | 3,394 | 0,765 | 3,391 | 0,090 | 5,113 | 0,783 |
| Molnupiravir | Nanchangmycin | RdRp/Tyrosine kinase | 30 | 1 | 3,524 | 0,006 | 3,489 | 0,805 | 5,260 | 0,224 |
| | | | 10 | 0,3 | 3,512 | 0,368 | 3,072 | 0,027 | 1,719 | 0,336 |
| Molnupiravir | Narasin | RdRp/Antibiotic | 10 | 1 | 3,085 | 0,395 | 3,116 | 0,520 | 3,842 | 0,115 |
| Molnupiravir | Nelfinavir | RdRp/Protease | 30 | 30 | 4,132 | 0,214 | 3,494 | 0,133 | 4,633 | 0,234 |
| | | | 10 | 10 | 2,932 | 0,408 | 3,182 | 0,047 | 3,810 | 0,740 |
| Molnupiravir | Niclosamide | RdRp/Multiple MOA | 20 | 10 | 3,190 | 0,180 | 2,812 | 0,661 | 3,236 | 0,368 |
| | | | 10 | 10 | 2,520 | 0,398 | 3,900 | 0,253 | 4,230 | 0,430 |
| Molnupiravir | Nitaxozanide | RdRp/Multiple MOA | 20 | 10 | 3,774 | 0,329 | 3,498 | 0,019 | 4,398 | 0,011 |
| | | | 10 | 30 | 3,012 | 0,066 | 3,265 | 0,110 | 3,806 | 0,094 |
| Molnupiravir | ONO-3307 | RdRp/Protease | 10 | 30 | 2,780 | 0,350 | 4,286 | 0,048 | 4,914 | 0,017 |

| | | | | | | | | | |
|---|---|---|---|---|---|---|---|---|---|
| Molnupiravir | PF-00835231 | RdRp/protease | 10 | 30 | 2,921 | 0,005 | 4,094 | 0,089 | 4,732 | 0,270 |
| | | | 10 | 5 | 3,128 | 0,082 | 3,370 | 0,144 | 4,096 | 0,468 |
| Molnupiravir | Remdesivir | RdRp/RdRp | 10 | 1,15 | 3,040 | 0,031 | 4,000 | 0,620 | 5,379 | 0,192 |
| | | | 10 | 0,36 | 2,322 | 0,948 | 2,775 | 0,416 | 3,086 | 0,790 |
| Molnupiravir | Ritonavir | RdRp/Protease | 10 | 10 | 2,619 | 0,567 | 3,148 | 0,054 | 3,515 | 0,398 |
| Molnupiravir | Samotilisib | RdRp/PI3K, mTOR | 10 | 1 | 0,939 | 0,295 | 1,580 | 0,317 | 1,476 | 0,247 |
| | | | 10 | 3 | 2,546 | 0,077 | 1,817 | 0,187 | 1,591 | 0,132 |
| Molnupiravir | TO-195 | RdRp/Protease | 10 | 30 | 2,945 | 0,496 | 3,906 | 0,366 | 4,572 | 0,377 |
| Nelfinavir | Camostat | Protease/TMPRSS2 | 30 | 30 | 3,424 | 0,132 | 4,235 | 0,411 | 4,792 | 0,051 |
| | | | 10 | 10 | 2,829 | 0,392 | 2,890 | 0,873 | 2,027 | 2,370 |
| Nelfinavir | Nafamostat | Protease/TMPRSS2 | 10 | 10 | 2,940 | 0,083 | 3,466 | 0,054 | 3,792 | 1,073 |
| Remdesivir | Nafamostat | RdRp/TMPRSS2 | 1,15 | 10 | 3,741 | 0,060 | 4,832 | 0,146 | 5,068 | 0,079 |
| | | | 3,3 | 30 | 3,476 | 0,077 | 4,545 | 0,031 | 5,199 | 0,473 |
| Ritonavir | Camostat | HIV protease/TMPRSS2 | 30 | 10 | 2,854 | 0,216 | 3,448 | 0,440 | 3,459 | 0,275 |

MoA: Mechanism of Action, AW: Apical Wash, IC: Intracellular, SD: Standard Deviation
